# Early morphogenetic patterns of protonemata and gametophores of *Physcomitrium patens* (Hedw.) Mitt

**DOI:** 10.1101/2024.07.22.604381

**Authors:** M.V.S. Raju, N.W. Ashton

## Abstract

In axenic culture, protonemata of *Physcomitrium* (formerly *Physcomitrella*) *patens* (Hedw.) Mitt. are comprised of chloronemata and caulonemata that can be distinguished morpho-structurally. On solid nutrient agar medium, a protonemal inoculum proliferates, initially generating primary chloronemata, and expands radially across the surface of the substratum forming approximately round colonies. The expansion of the colonies derives mainly from centrifugal elongation of newly formed caulonemata, which arise by differentiation of primary chloronemal apical cells in the central region. Most commonly, oblique cross walls separate cells of the uniseriate caulonemal filaments. Each oblique wall organises a smaller and a larger angle with the outer wall of a tubular caulonemal cell. The smaller angle may lie in four different positions, vertically upward or downward, and laterally, to the right or left, parallel to the surface of the solid medium. Chloronemal filaments, which contain cells separated by transverse cross walls, develop from side branch initials (SBIs) at or near the smaller angle at the distal end of caulonemal cells. Like the smaller angle of the oblique cross walls in caulonemata, these secondary chloronemata also show four different orientations, vertically, upward or downward, and laterally, right or left. A strong inverse correlation exists between the placement of SBIs and secondary chloronemata and the direction of caulonemal tip growth. If the caulonemal tips curve downward into the medium, chloronemata are situated vertically on the upper side of caulonemata; similarly, if they turn upward, chloronemata are situated on the lower side. A similar correlation exists when caulonemal tips curve to the right or left on the surface of solid medium, in which case laterally directed chloronemata grow in the opposite direction, i.e. on the outside of the curve. The procumbent caulonemata with their associated protruding secondary chloronemal branches are reminiscent of the heterotrichous branching habit of filaments of some green algae. Gametophore buds develop from a minority of caulonemal SBIs and also occasionally from chloronemata; a stalk cell, which resembles a chloronemal cell, supports each bud. Buds develop into leafy gametophores that are arranged in a fairy ring. We discuss the relative merits of several models that might explain the temporally and spatially regulated clustering of buds resulting in the formation of a fairy ring.

## Introduction

Since the seminal paper by Paulinus Engel (1968), *Physcomitrella* (recently renamed *Physcomitrium*) *patens* has become a popular model plant, which has received more than 20,000 citations (Duckett 2018). Quite a lot has been published about the hormonal regulation of its morphogenesis (e.g. Ashton et al. 1979, Decker et al. 2006, Proust et al. 2011, Thelander et al. 2018) and the tropic responses of its protonemata and gametophore shoots (e.g. Knight & Cove 1989). Its use as a model system for studying plant development received a significant boost at the end of the last century from the first of many reports of successful gene targeting experiments (e.g. Girke et al., 1998, Strepp et al., 1998). Realistic computer modelling of its protonemal development has also been achieved (Ashton et al. 1993, Fracchia & Ashton, 1995), facilitated by the ease of observation of cell lineages in the branching system of filaments that comprises the protonema of *P. patens.* Therefore, it is surprising that basic information about the morphogenetic patterning of *P. patens* protonemata and gametophore bud initiation remains sparse. Perhaps the most informative data are those provided by McClelland (1987, 1988), Russell (1993) and recently by Antonishyn et al. (2024). McClelland summarises quantitative data describing patterns of protonemal branching in 21 day old colonies. In brief, he observes that 1) the development of SBIs on caulonemata is restricted to the second subapical and older cells; 2) most such initials become chloronemata (85-90%); 3) a minority of initials develop into caulonemata (5-6%) or buds (<1%); 4) a maximum of one bud is produced on each caulonema. Russell extended the observations of McClelland especially with respect to the formation of a second SBI on older caulonemal subapical cells and also the clustering of buds on individual caulonemata. Antonishyn et al. provide a detailed account of the spiral morphology of *P. patens* colonies and possible mechanisms responsible for the curvature of radiating caulonemata. In an earlier paper, Bopp (1959) reported similar qualitative findings about caulonemal curvature in *Funaria hygrometrica*, a close relative of *P. patens*, both belonging to the family Funariaceae. He also noted gametophores were initially formed in a Hexenringe or fairy ring (Bopp 1952), which we have observed in *P. patens* (Ashton 1974). Herein we have focused on compensating for omissions in information about the early morphogenetic patterns displayed by *P. patens* gametophytes by presenting the results obtained from observations made on laboratory cultures grown from vegetative inocula during a 25 day period using standardised conditions and standard nutrient agar medium. Nitrate was chosen as sole source of nitrogen since other nitrogen sources significantly alter morphogenetic patterning, e.g. ammonium increases substantially the production of chloronemata, while some other changes to the medium have the opposite effect, e.g. omission of phosphate dramatically reduces the production of chloronemata (McClelland 1987, 1988).

We hope that our findings will contribute significantly to a description of normal morphogenesis in *P. patens*, which can be used as a meaningful baseline for comparison to and interpretation of altered developmental patterns induced by experimental manipulation of the environment and/or genome of this moss. Also, these data should lead to further improvement of computer models of *P. patens* morphogenesis and bring closer realisation of one of the major goals of this approach, namely to generate models of sufficient quality that, when manipulated, thev will provide reliable predictions about the behaviour of the real plant when it is treated in an analogous way.

## Materials and methods

Protonemal inocula were taken from stock cultures of wild type *P. patens* derived from a single spore isolated from nature in Gransden Wood, Huntingdonshire, U.K. in 1962 by H.L.K. Whitehouse. Since then, this isolate has been maintained in axenic culture by repetitive, serial somatic cloning of gametophytes, interspersed occasionally and sporadically during its early history with passages through the complete sexual life cycle. Gametophytes were grown on standard ABC culture medium (Knight et al. 1988) solidified with 1.5% (w/v) agar (Sigma, MO, USA) in plastic Petri dishes. The thickness of the nutrient agar medium in the Petri plates was about 5 mm. Cultures were incubated at 22-25°C under continuous light supplied by cool white fluorescent tubes (Westinghouse, Regina, SK, Canada). The Petri plates were covered with one layer of clear (Roscolux, No. 114, Hamburg frost; MacPhon Industries, Calgary, AB, Canada) resin filter (to reduce the rate of evaporative water loss from the culture medium). Photon flux of PAR, measured with a quantum photometer (model LI-185 A, Li-cor, Lincoln, NEB, USA) connected to a quantum sensor (model LI-190S), was 66-107 µmol m^-2^ s^-1^ at the surface of the medium.

Twenty five small vegetative inocula were placed on the surface of solid growth medium in five 9 cm diameter Petri plates, five inocula per plate. In each plate, a central inoculum was surrounded by four equidistant peripheral inocula. From and including the third day after inoculation, colonies derived from these inocula were observed every second day with a stereoscope and morphological details recorded. Observations were terminated after 25 days. The growth of each colony was assessed at regular intervals by measuring the widest horizontal dimension of the approximately round colonies. After the first five or six days, this was typically the distance between the apical cells of two caulonemata radiating in opposite directions from the central region of primary chloronemata. The number of gametophores (buds and leafy shoots) in each colony was also counted at regular times and recorded. Random samples of protonemal filaments from colonies of comparable age cultured in parallel with those described above were isolated and examined with a compound microscope fitted with a camera in order to observe and record cellular and morphological details.

### Mathematical analysis and statistics

Numerical data were fitted to curves using TableCurve (Jandel Scientific, San Rafael, CA, USA), which also provided the following goodness of fit measurements: Coef of Det (Coefficient of Determination) r^2^, DOF (Degrees Of Freedom) adjusted r^2^, Fit Std Error (Fit Standard Error) and F-statistic. The closer the value of r2 to 1.0, the better is the fit. The Fit Std Error is the least squares error of fit and the closer the value to zero, the better the fit. The higher the F-statistic, the better an equation models the data. Averages and standard deviations were calculated using Microsoft Excel.

## Results

### The general pattern of protonemal development

Two to three days after being inoculated on to fresh medium, cells in an inoculum could be seen to be regenerating. They proliferated to produce a central region of short, branching chloronemal filaments (defined herein as primary chloronemata, since cytologically and behaviourally they are identical to the primary chloronemata from germinated spores) on the surface of the solid medium. The cells of the filaments had hyaline walls and contained round, dark green chloroplasts. After approximately three more days, many caulonemata emerged from the central chloronemal region and continued growing horizontally and centrifugally on or close to the surface of the medium. The elongating caulonemal filaments produced branches most of which were positively phototropic chloronemata that ultimately grew upwards. After approximately 20 to 25 days, the central area of chloronemata began to degenerate while the rest of the colony continued expanding centrifugally (**Fig.1**).

**Fig. 1.**
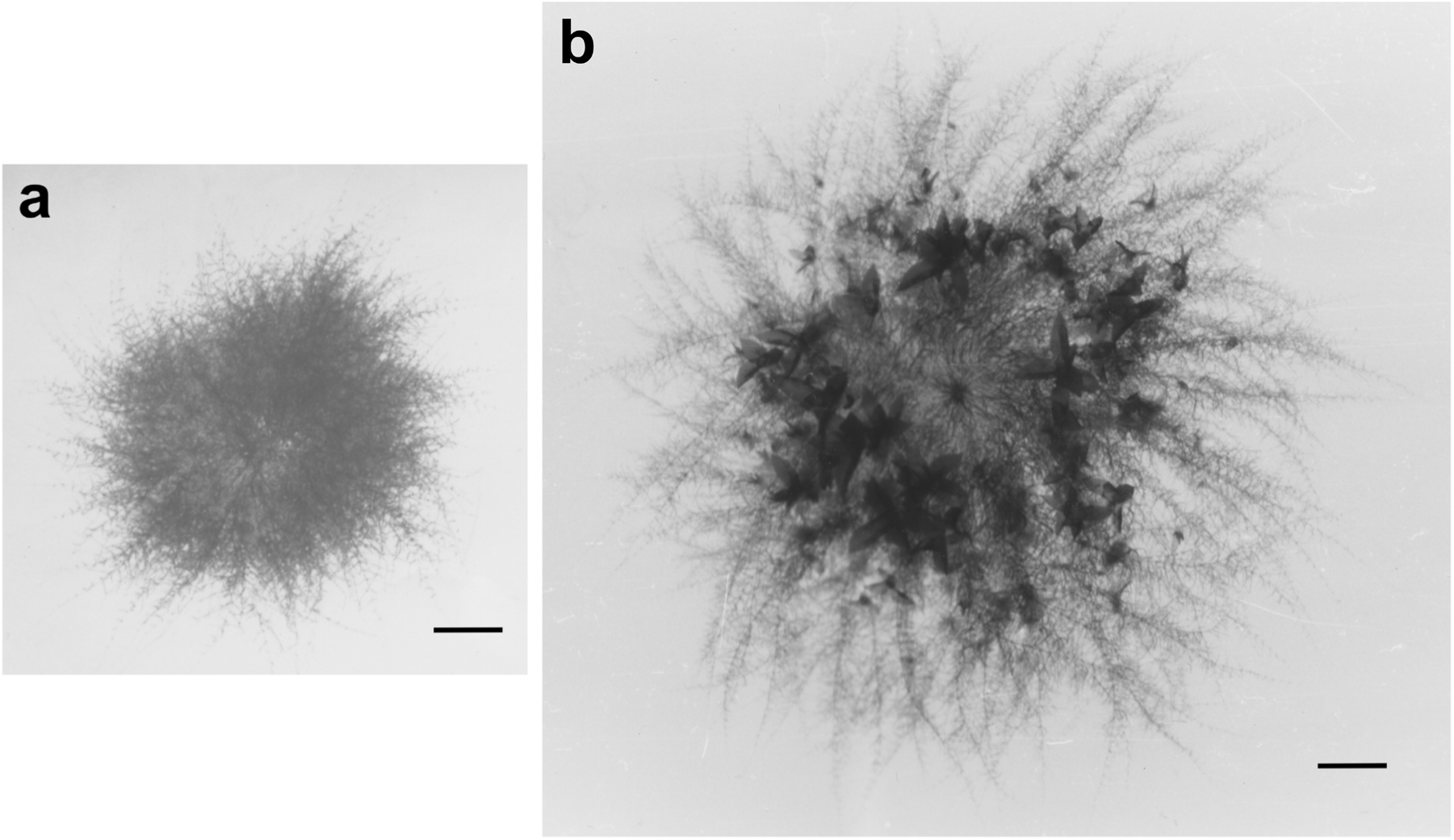
Gametophytes of *P. patens*. ***(a)*** Two week old colony comprised of central primary chloronemata from which centrifugally radiating, procumbent primary caulonemata with protruding secondary chloronemata have differentiated. Gametophore buds are present near the periphery of the colony but have not grown and developed enough to be visible at this magnification. ***(b)*** Three week old colony with a fairy ring of leafy gametophores and peripheral caulonemata with secondary chloronemata. The relative sizes of the gametophytic images reflect the relative sizes of the real gametophytes. Scale bars = 1 mm.

### Primary chloronemata produced by regeneration of cells of an inoculum

Inocula generally consisted of a few chloronemal and caulonemal filaments but occasionally included a bud or leafy gametophore. On fresh medium, primary chloronemata regenerated from any cell or tissue type present in an inoculum. Chloronemal apical cells usually originated as branch initials on protonemal cells but also arose from individual surface cells of buds, gametophore stems and leaves. The new chloronemal apical cells had transverse (perpendicular to the long axis of the cells) cross walls at their proximal end; they elongated and divided, forming uniseriate filaments of cells with transverse dividing walls (**Fig. 2a**). Subapical cells of these new chloronemal filaments themselves produced initials, which also became chloronemal apical cells by the formation of a transverse cross wall parallel to the long axis of the cell upon which the initial formed. Thus, most filaments in the central chloronemal region of young colonies were branched.

**Fig. 2.**
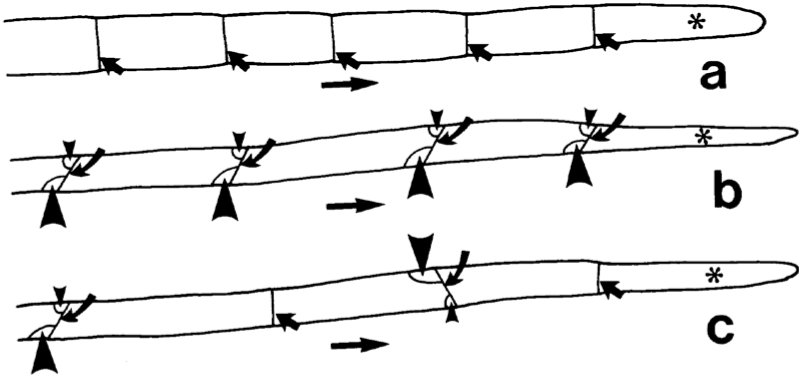
Schematic line diagrams of the few most distal cells of young protonemata. ***(a)*** A chloronemal filament with wider cells and transverse cross walls. The cross wall at the distal end of each cell organises a 90° angle with the side wall. ***(b)***

A tapered caulonemal filament with narrower cells separated by oblique cross walls. Each oblique cross wall organises with the side wall one larger and an oppositely placed smaller angle. ***(c)*** A tapered caulonemal filament with narrower cells separated by both transverse and oblique cross walls. The horizontal arrows indicate the direction of filament growth. Cells with an asterisk (*) are the apical cells of filaments. Short straight arrows point at transverse cross walls; curved arrows indicate oblique cross walls; larger arrowheads indicate larger angles between cell walls; smaller arrowheads point at smaller angles.

### Caulonemata

By approximately five or six days after inoculation, some of the chloronemal apical cells had differentiated into caulonemal apical cells, which elongated and divided producing uniseriate primary caulonemata (**Figs. 2b, c, 3a, b**, **4**) that emerged from the periphery of the central area of primary chloronemata. Typically, caulonemal apical cells possessed an oblique cross wall at the proximal end and the caulonemal filaments derived from them by repeated cell division consisted of cells whose dividing walls were also predominantly oblique. Each oblique cross wall organised a smaller and a larger angle with the longitudinal wall of a tubular caulonema. The larger angle at the distal end of caulonemal subapical cells varied between 105° and 150° with a mean value of 136.35° and standard deviation of ± 12.64° (N=25). The smaller angle was observed to lie mainly in four positions in two planes: in the vertical plane, on the upper or lower side of a caulonema; in the horizontal plane, on the left or right side of a filament. Occasionally a transverse cross wall was laid down during division of a caulonemal apical cell. Thus, a few transverse dividing walls were seen between subapical cells in caulonemata (**Figs. 2c, 3a, b, 4a (panel A), 4d (panel C), 4e** (panel D)).

**Fig. 3.**
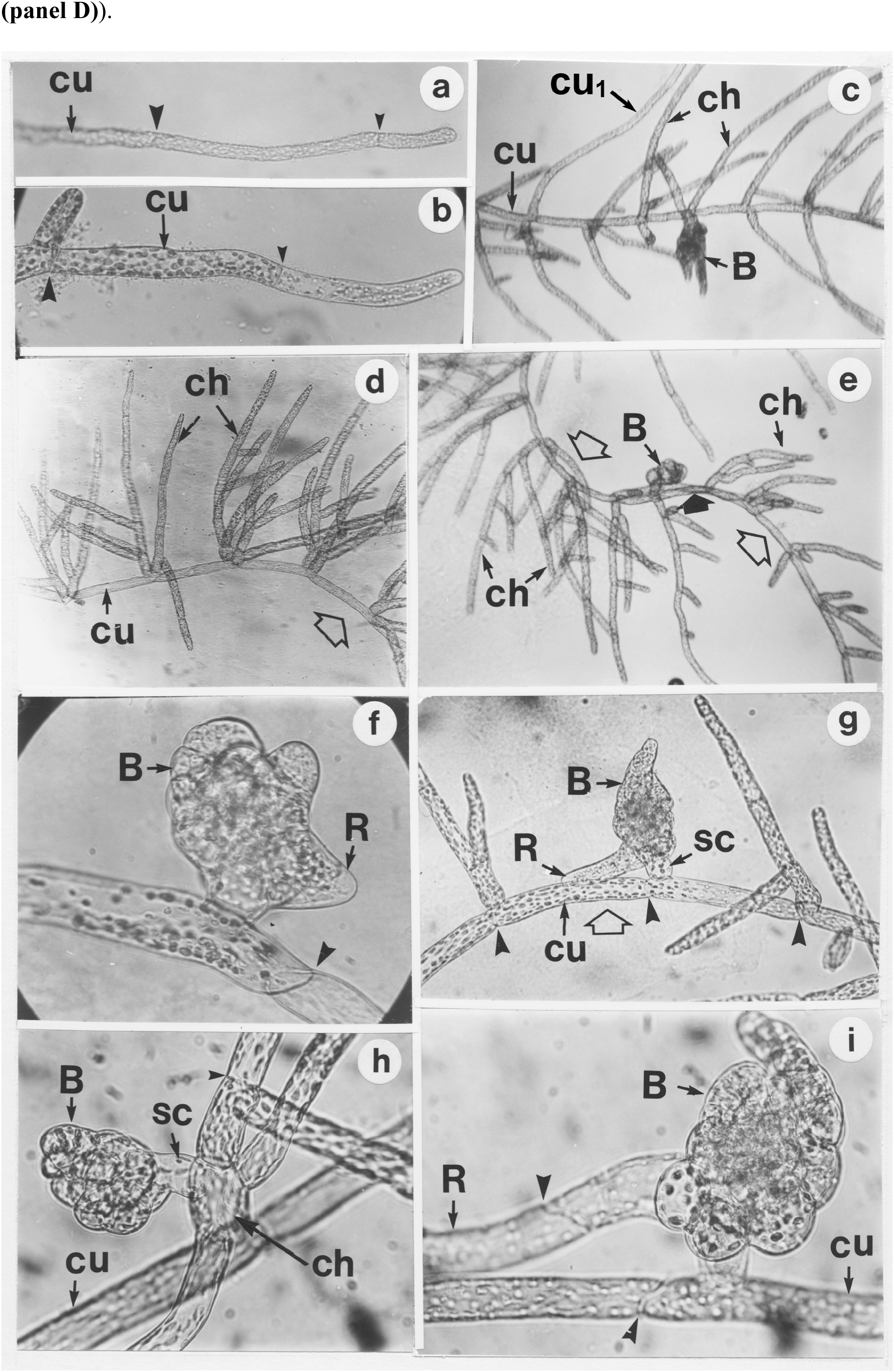
Growth and morphology of caulonemata, secondary chloronemata and gametophore buds. *(a)* Distal tip of a thin caulonema with a transverse and an oblique cross wall, x 350. *(b)* A thick caulonema with a chloronemal side branch originating near the smaller angle formed by the oblique wall at the distal end of the caulonemal subapical cell, x 220. *(c)* A proximal section of a caulonema with most subapical cells possessing two secondary chloronemal side branches at their distal end. A single bud is present at the distal end of one of the caulonemal subapical cells and a secondary caulonema has arisen from another, x 125. *(d)* A proximal section of a procumbent caulonema with two protruding and branching secondary chloronemata arising from each subapical cell forming a V-shaped structure. The caulonema is curved rightwards (clockwise) with side branches on the outside of the curve. x 125. *(e)* A caulonema with alternating clockwise (at its distal end, on the right) and anticlockwise (at its proximal end, on the left) curvature. In both regions, the chloronemal branches arise on the outside of the curve, x 140. *(f)* A gametophore bud situated at the distal end of a caulonemal subapical cell adjacent to the smaller angle subtended by the cross wall, x 350. *(g)* A distal section of a caulonema with with a single, branching chloronema or a bud arising from each subapical cell. The chloronemata have a first order side branch developing from their basal cell forming an asymmetric Y-shaped structure and are situated on the outside of the anticlockwise curve of the curved caulonema. The bud, with a rhizoid from its base, is supported by a stalk cell, x 330. *(h)* A lateral bud on the basal cell of a chloronemal side branch of a caulonemal subapical cell, x 580. *(i)* An older bud, with a rhizoid at its base, arising directly from the distal end of a caulonemal subapical cell x 750. B = gametophore bud; ch = chloronema; cu = caulonema; cu_1_ = secondary caulonema; R= rhizoid; sc= stalk cell of gametophore bud. Small arrowheads indicate transverse cross walls and large arrowheads point to oblique cross walls; unfilled wide arrows point to the curvature in caulonemata; a solid wide arrow indicates an inflection in the curvature of a caulonemal filament (anticlockwise to clockwise) and a concomitant change in the side of origin and growth direction of secondary chloronemata on the caulonemal subapical cells. The walls of caulonemal apical cells and of young subapical cells were hyaline in contrast to those of older subapical cells, which accumulated a red-brown pigment. Wall pigmentation appeared in caulonemata when the colonies were between 10 and 15 days old, during the time gametophores were formed, and was more pronounced in caulonemal cells proximal to a gametophore than in cells distal to it.

**Fig. 4.**
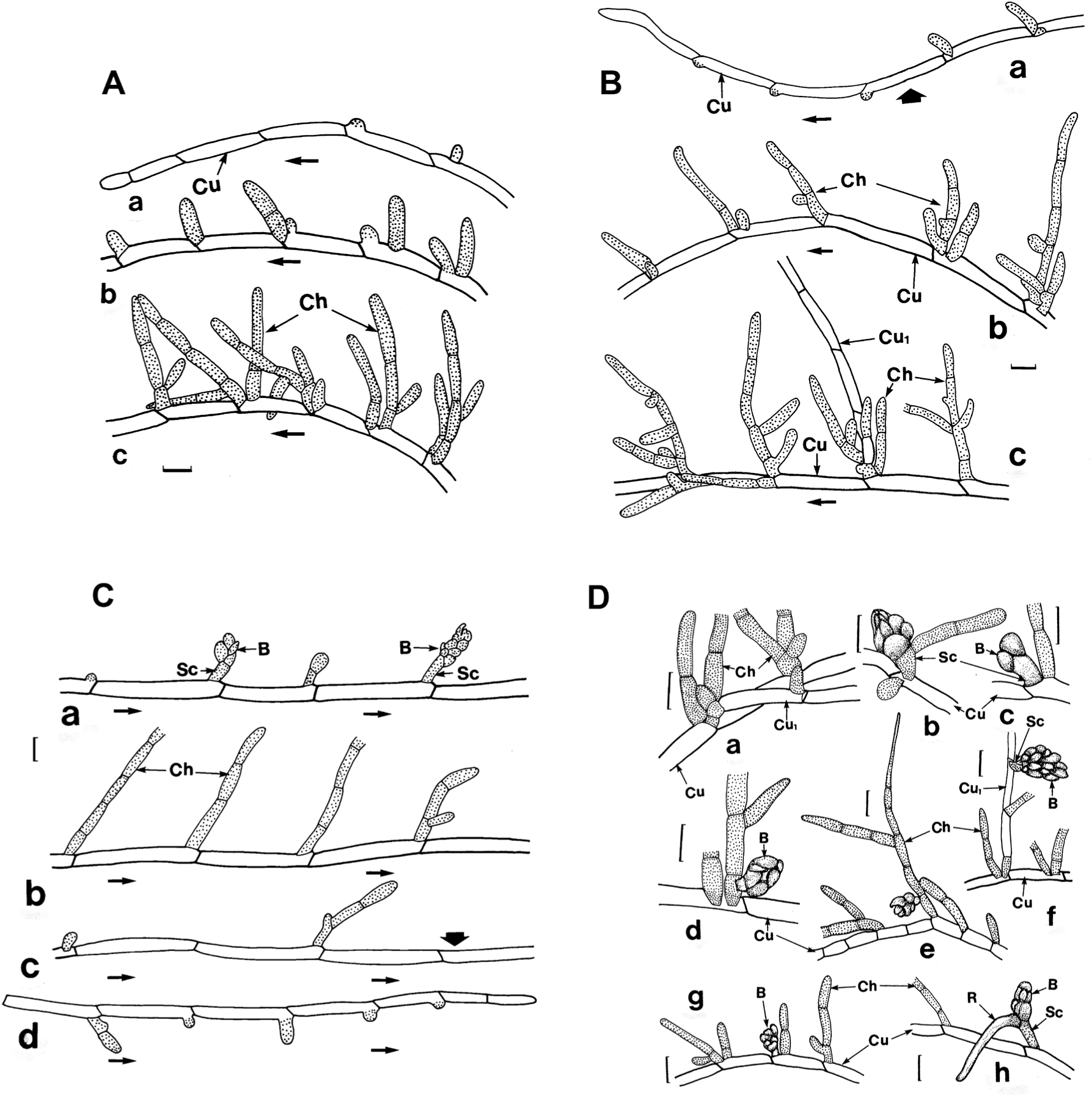
Drawings illustrating the variety and features of caulonemata and secondary structures (papillae, SBIs, secondary chloronemata, secondary caulonemata and gametophore buds) that develop from their subapical cells. **Panel A.** Three consecutive, contiguous distal sections ***(a, b, c)***of a caulonema (not stippled). Arranging the segments end-to-end in sequence, this caulonema possesses anticlockwise curvature with protruding SBIs and secondary chloronemata on the outside of the curve. All the cross walls betweeen caulonemal subapical cells are oblique except for the transverse wall between the apical cell and the first subapical cell. Horizontal arrows point to the distal end (apex) of the filament. **Panel B.** A caulonema, showing distal (top left) and proximal (bottom right) regions arranged in sequence, with papillae on the first three subapical cells, SBIs and secondary chloronemata (all stippled) on more proximal cells, and a single secondary caulonema (not stippled). Arranging the caulonemal sections ***(a, b, c)*** end-to-end, in sequence, reveals a change in profile from clockwise curvature ***(a, distal end)*** to anticlockwise curvature ***(a, proximal end, and b)*** to straight ***(c)***. A vertical, filled wide arrow indicates the point of inflection in the curvature. Secondary structures arise on the outside of the curve of curved regions of the caulonema. **Panel C.** Four sections in sequence ***(a, proximal, to d, distal)*** of a straight caulonema. The caulonema had its origin in a young leafy gametophore. Papillae, SBIs, secondary chloronemata and young gametophore buds (all stippled) arise at the distal end of a caulonemal subapical cell next to the acute angle formed by each oblique cross wall. The oblique cross walls of contiguous caulonemal subapical cells tend to be aligned and thus all the structures that arise from them tend to do so on the same side of the filament. The vertical, filled wide arrow indicates a transition in the side of origin and growth direction of secondary structures on the caulonema. **Panel D.** Segments of procumbent caulonemata and associated protruding secondary chloronemata and buds. ***(a)*** Chloronemata at the distal end of caulonemal cells near the smaller angle subtended by each oblique cross wall. One of the chloronemal filaments has differentiated into a first order caulonemal branch. ***(b)*** A segment of a primary caulonema with chloronemal side branches, one of which has produced a gametophore bud from its basal cell. ***(c)*** A gametophore bud with its stalk cell situated at the distal end of a caulonemal subapical cell. ***(d)*** A segment of a caulonema with two chloronemal side branches, one of which has a lateral gametophore bud on its basal cell. ***(e)*** A caulonema with rare transverse cross walls and chloronemal side branches, one of which has a lateral gametophore bud on its basal cell. ***(f)*** A caulonemal segment with two chloronemata in an asymmetric V-shaped arrangement at the distal end of each caulonemal subapical cell. One of the chloronemal branches has differentiated into a secondary caulonema bearing a chloronema and a gametophore bud with a stalk cell. ***(g)*** A caulonema with secondary chloronemata and a gametophore bud. ***(h)*** A caulonemal filament with a gametophore bud, which has a stalk cell and a rhizoid. Horizontal arrows indicate the direction of caulonemal growth. B = gametophore bud, Ch = chloronema, Cu = caulonema, Cu_1_ = a first order, secondary caulonema developing from the basal cell of a chloronemal side branch, R = rhizoid, Sc = stalk cell of gametophore bud. Scale bars = 50 µm.

The apical cell of caulonemata contained many small, round amyloplasts towards its distal end but proximal to the apical dome, which appeared to be free of organelles. More distal subapical cells tended to contain relatively few chloroplasts while more proximal subapical cells contained abundant, elongated, spindle-shaped chloroplasts, unlike the abundant, large round chloroplasts that filled both primary and secondary chloronemal apical and subapical cells.

Many of the indeterminate caulonemata were procumbent and grew across the surface of the solid medium; some grew as subsurface filaments within the medium. All caulonemal subapical cells had the potential to produce SBIs that developed predominantly into secondary chloronemal branches (**Fig. 4**).

Branch initials arose from caulonemal and chloronemal subapical cells at the distal end of the cells, indicating polarity in both cases (**Figs. 3b, 4**). In contrast to chloronemal cells, where only one branch per cell developed, the majority of caulonemal cells produced sequentially two separate branches. Thus, young caulonemal subapical cells had one SBI or side branch; older cells were typically characterised by two chloronemal side branches, the shorter more recent of which arose near the longer first branch at the distal end of the cell, creating an asymmetric V-shaped arrangement of side branches emanating from the caulonemal subapical cell (**Figs. 3c-e**, **4b, c (panel A), 4b (panel B), 4f (panel D)**). Occasionally one of the side branches developed into a gametophore bud instead of a secondary chloronema (**Fig. 3e-g**, **4a (panel C), 4c, g, h (panel D)**). The development of two separate bud-bearing branches from the same caulonemal subapical cell was not observed. A minority of the older, more proximal caulonemal subapical cells had only a single secondary chloronema, or a single bud, or a single SBI that had failed to differentiate, or nothing (**Fig. 4 (panel C)**).

SBIs, which differentiated predominantly into secondary chloronemata but also into secondary caulonemata and gametophore buds or occasionally remained dormant, were formed by division of the second or third or older subapical cell of each caulonema. Initially a single SBI was formed at the distal end of these caulonemal subapical cells at or near the smaller angle made by the oblique cross walls that characterise caulonemata (**Figs. 4a**, **b (panel A), 4a (panel B), 4c, d (panel C)**). These smaller angles, depending on the orientation of the cross walls, were aligned in a row along the upper, lower, left or right side of each tubular caulonemal filament. This resulted in the corresponding alignment of SBIs and their derivative structures (secondary chronemata, secondary caulonemata and buds), i.e. in a row on the upper, lower, left or right side of a caulonema. There was also a strong correlation between the direction of growth of caulonemal apical cells and the orientation of the oblique cross walls between the subapical cells. If the apical cell grew downwards, i.e. towards or into the medium, the smaller angles of the oblique cross walls occurred on the upper side of the caulonema (**Fig. 5a**). If the apical cell grew upwards, i.e. away from the medium, the smaller angles of the cross walls were situated on the lower side (**Figs. 5b**, **c**). In instances where the apical cell grew to the left or right, resulting in an anticlockwise or clockwise curvature respectively of the entire caulonema, the smaller angles were arranged on the right or left, i.e. laterally on the opposite side of the curve, of the tubular filament (**Fig. 5d**). Occasionally, after several cell divisisions, a caulonemal apical cell changed direction, e.g. from growing upwards to downwards, causing the caulonema to assume an undulating appearance (**Fig. 5a-c**). These morphogeneic patterns of caulonemata with associated protruding secondary branches were observed most readily in 15 to 25 day old colonies in which the caulonemata had elongated considerably.

**Fig. 5.**
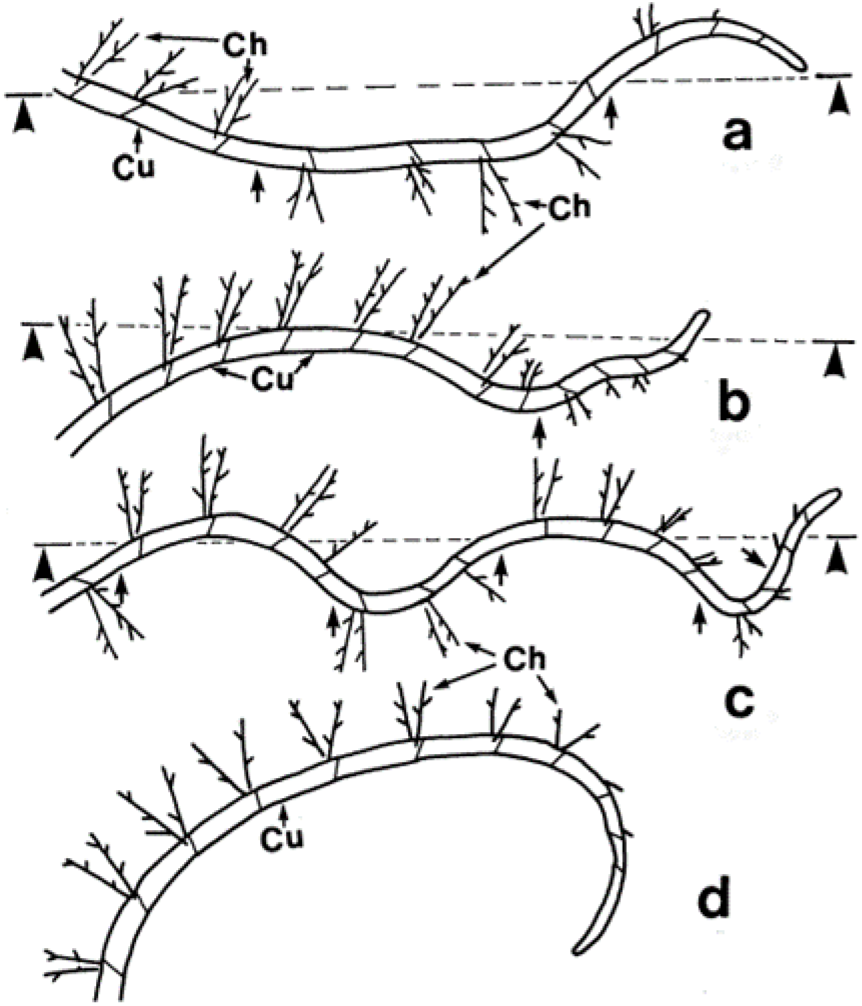
Schematic line drawings of protonemata on solid medium, illustrating several morphogenetic patterns of caulonemal and secondary chloronemal growth. Each caulonema shows the proximal end to the left and the distal end to the right. The lines indicated by arrow heads represent the plane of the horizontal air/solid medium interface. ***(a, b, c)*** In these examples, the direction of growth of the caulonemal apical cell alternates in the vertical plane between downwards and upwards, resulting in caulonemata with an undulating profile and subsurface and subaerial sections. Arrows indicate inflection points in caulonemal curvature and changes in the sideness of secondary chloronemata and in the orientation of oblique cross walls between caulonemal subapical cells. ***(d)*** View from above of a caulonema with chloronemata unilaterally disposed on the outside of the clockwise curvature of the filament. A caulonema may have anticlockwise curvature (not presented here), also with chloronemata emanating from the outside of the curve. Clockwise curvature on the surface of solid medium is the predominant form of caulonemal curvature in *P. patens* illuminated from above. Ch *=* chloronema, Cu *=* caulonema.

Caulonemal apical cells had five main origins. Primary caulonemata differentiated from primary chloronemal apical cells (**Fig. 3a**) of the central chloronemal region five or six days after inoculation. Secondary caulonemata developed about 12-15 days after inoculation from SBIs on pre-existing caulonemata (**Fig. 3c**), branch initials formed on basal cells of secondary chloronemata (**Fig. 4c (panel B)**), apical cells of secondary chloronemata and single cells at the base of a bud.

The differentiation of primary caulonemata at about day five or six was reflected in the increased width of the colonies (**Fig. 6a**). Continued expansion of the colonies resulted predominantly from the centrifugal growth of primary caulonemata and of secondary caulonemata that appeared about 12 to 15 days after the start of the experiment. Some of the secondary caulonemata grew in different directions, filling in spaces between the previously formed radial primary caulonemata and contributing to the density as well as expansion of the protonemal colonies. Such growth changes sometimes altered the original circular morphology of colonies making them more irregular in shape. Changes in colony morphology were also caused by caulonemata developing from the base of young gametophores.

**Fig. 6.**
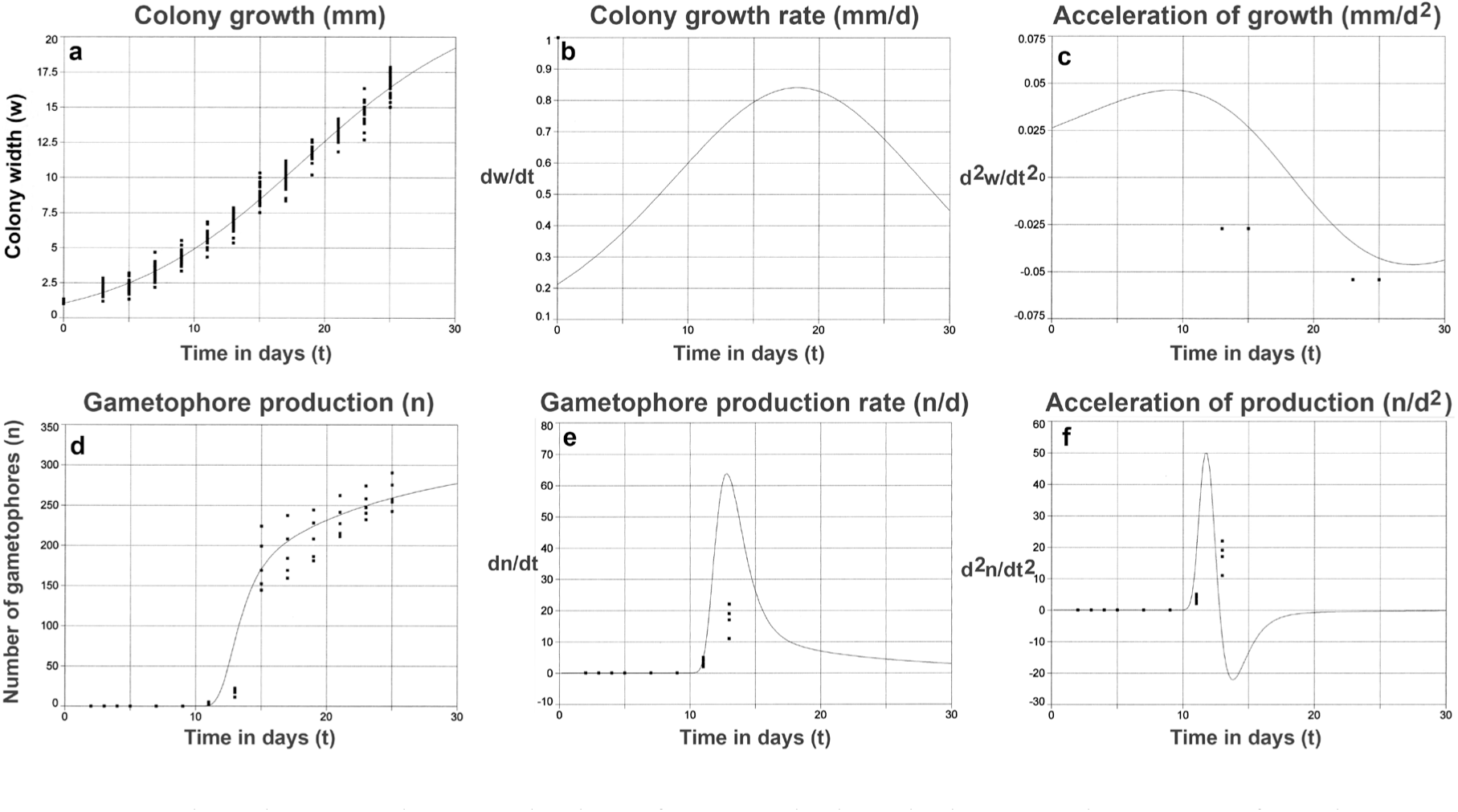
Growth and gametophore production of gametophytic colonies over the course of 25 days. ***(a)*** Maximum width of colonies (n = 25) in mm plotted against time in days, ***(b)*** the first differential of the growth data, which gives the instantaneous growth rate versus time, ***(c)*** the second differential of the growth data, which provides the instantaneous acceleration of growth versus time. ***(d)*** Number of gametophores/Petri dish (n = 5 with 5 colonies/dish, 25 colonies in total) plotted against time, ***(e)*** the first differential of the gametophore numbers data, which gives the instantaneous gametophore production rate versus time, ***(f)*** the second differential of the production data, which provides the instantaneous acceleration of gametophore production versus time.

Over the course of the experiment, the size of colonies conformed closely to a sigmoid growth curve defined by the function, y=a+b/(1+exp(-(x-c)/d)), where y=maximal colony width (w in mm), x=time (t in d), a=0.55904716, b=23.485281, c=18.334728, d=6.986366 (**Fig. 6a**). r^2^=0.984188381, DF Adj r^2^=0.983890736, FitStdErr=0.638631444, Fstat=6660.17541. Differentiation and redifferentiation of the data gave plots of the first differential, i.e. the instantaneous growth rate (dw/dt) (**Fig. 6b**) and the second differential, i.e. the instantaneous rate of change of the growth rate (d^2^w/dt^2^) (**Fig. 6c**) respectively. We refer to the latter (d^2^w/dt^2^) loosely as acceleration since there is no scalar term corresponding to the vector, acceleration, in the English language. Acceleration of the growth rate was maximal at day nine and the growth rate increased to a maximum at day 18 or 19 and thereafter declined.

### Gametophore buds

Buds, which developed into leafy gametophores, usually arose from caulonemal cells (**Figs. 3f, g, i, 4a (panel C), 4b, c, g, h (panel D)**). A small protuberance or papilla appeared at the distal end of a caulonemal subapical cell (**Figs. 4a (panel A), 4a (panel B), 4d (panel C)**) and became separated from it forming a SBI. The initial frequently became a chloronemal branch as described below. Alternatively, it divided producing a short filament of two cells, the distal bud initial and the proximal stalk cell. The bud initial divided further to develop into a bud and eventually a leafy gametophore. The proximal cell became a long or short stalk cell subtending the bud (**Figs. 3f, g, i, 4a (panel C), 4b, c, f-h (panel D)**). The stalk cell resembled morphologically a chloronemal cell, containing round green chloroplasts. Such a stalked bud sometimes developed on the basal cell of a chloronemal branch of a caulonema (**Figs. 3h, 4d, e (panel D)**; also observed by Thelander 2018). The number of buds produced on a mature caulonema varied from one to three (**Fig. 4a (panel C**). No buds were observed on the first order branches of chloronemata.

Bud initials began appearing at about day11. Over the course of the experiment, the number of gametophores conformed closely to a curve defined by the function, lny=a+b/x^1.5^+ce^-x^, where y=number of gametophores (n), x=time (t in d), a=5.8392412, b=-35.335469, c=-302530.36 (**Fig. 6d**). r^2^=0.954997545, DF Adj r^2^=0.952951979, FitStdErr=24.0279525, Fstat=710.903835. Differentiation and redifferentiation of the data gave plots of the instantaneous rate of gametophore production (dn/dt) (**Fig. 6e**) and the instantaneous rate of change of dn /dt (d^2^n/dt^2^) (**Fig. 6f**) respectively. Acceleration (d^2^n/dt^2^) was maximal at day 12 and the rate of bud production (dn/dt) increased to a maximum at day 13, then declined sharply until day 15 and thereafter decreased more slowly.

### Rhizoids

In 23 to 25 day old cultures, colourless rhizoids had emerged from the base of buds and young leafy gametophores (**Figs. 3f, g, i, 4h (panel D)**) and grew downwards into the medium. In other respects, they closely resembled caulonemata and their cells possessed oblique cross walls. The larger angle of the distal oblique wall of rhizoidal subapical cells varied between 112° and 150° with a mean value of 132° and SD ± 10° (N=25). These values are similar to those for caulonemal cells.

The uniseriate filaments tapered towards the distal end and were generally unbranched and lacked papillae and SBIs. Proximal, i.e. closer to the base of gametophores, subapical cells tended to contain more abundant chloroplasts, which in older rhizoids were spindle-shaped, than distal subapical cells and the apical cell. The cell walls of rhizoids of older leafy gametophores turned red-brown.

### Secondary chloronemata derived from side branch initials on caulonemata

Each secondary chloronema was derived from a small outgrowth or papilla at the distal end of a caulonemal subapical cell that divided to form two cells. The formation of a cell wall separating the papilla from the subtending caulonemal cell created a SBI (**Figs. 4a**, **b (panel A), 4a, b (panel B), 4c, d (panel C)**). In most cases, the single-celled SBI was transformed into an apical cell, which divided several more times forming a short chloronemal filament of determinate growth, which was comprised of cells with thin hyaline walls separated by transverse, i.e. perpendicular to the long axis of the cell, cross walls (**Figs. 4b**, **c (panel A), 4b, c (panel B), 4b-d (panel C)**). Subapical cells of these determinate filaments divided once at their distal end giving rise to first order side branches, most of which were short and also displayed determinate growth (chloronemata) producing an asymmetrical Y-shaped structure (**Figs. 3g, 4c (panel A), 4b, c (panel B), 4b, c (panel C), 4g (panel D)**); a few, however, were characterised by indeterminate growth (caulonemata). No second order branches were observed on the chloronemal filaments during the experimental period. Occasionally a first order branch, especially one emanating from the basal-most cell of a chloronemal filament, displayed indeterminate growth (a caulonema) (**Figs. 4c (panel B), 4a (panel D)**). Also, some basal cells of chloronemata branched to produce a gametophore bud (**Figs. 3h, 4d, e (panel D)**). Seconday chloronemata were arranged contiguously along caulonemata or in groups intercalated with regions of their absence on the same caulonema.

Initially the disposition of secondary chloronemata on caulonemata conformed to four different patterns. They grew upwards (away from the surface of the medium), downwards (towards or into the medium) (**Figs. 5a-c**), or laterally (parallel to the surface of the medium) leftwards (anticlockwise) or rightwards (clockwise) (**Fig. 5d**).

## Discussion

The protonemata of mosses often comprise two interconvertible types, chloronemata and caulonemata Cove 1992, Vidali and Bezanilla 2012). The differentiation of one type into the other is a progressive process rather than an abrupt transition (herein and Jang and Dolan 2011, Thelander 2018). Thus, for example, acquisition of a polarised distribution of intracellular organelles and of an oblique proximal cross wall by a caulonemal apical cell differentiating from a chloronemal apical cell occurs gradually and the differentiation of a secondary chloronema from a caulonemal subapical cell occurs via a SBI. Also, the two types of protonemata are less readily distinguishable in some species than in others (Allsopp and Mitra 1956, 1958). Therefore, recognition of chloronemata and caulonemata should be based not on one character but on as many characters as possible, as has been done by Bopp (1961, 1990) and others, e.g. Duckett et al. (1998). Two characters relating to chloronemal and caulonemal cells highlighted by Duckett et al. (1998) are worthy of further comment, namely the mode of extension of apical cells and cell polarity. While it is generally claimed that both chloronemal and caulonemal apical cells elongate by tip growth (e.g. Vidali and Bezanilla 2012), Duckett et al. (1998) provide strong arguments that chloronemal apical cells do not and instead elongate by intercalary wall expansion. If true, the differentiation of a chloronemal apical cell into a caulonemal apical cell must involve complex reorganisation of the internal structure of the cell, as shown relatively recently by Jaeger and Moody (2021), that can only be achieved progressively over the course of several cell cycles as we have observed. Duckett et al. (1998) also contrast the polar versus non-polar distribution of organelles in caulonemal and chloronemal cells respectively. While we agree with this observation, both caulonemal and chloronemal subapical cells display polar behaviour when they divivide forming a side branch or bud. When this happens, the derivative structure always arises at the distal end of the cell. However, there is a difference in the possible positions of the structure with respect to the short axis of the cell. In caulonemal cells, it is confined to a location at or near the smaller angle formed by an oblique cross wall; in chloronemal cells, it can occur at any point around the circumference of the tubular cell since the transverse cross wall is consistently perpendicular to the long axis of the cell. Two morphogenetic patterns that occur during the gametophytic phase of the life cycle of *P. patens* were of particular interest to us in this study.

Procumbent indeterminate primary caulonemata with protruding determinate chloronemal side branches (secondary chloronemata) comprise a heterotrichous system of peripheral branching filaments radiating acropetally from primary chloronemata near the centre of the colony (**Figs. 3d, 4**). While SBIs formed on caulonemal subapical cells differentiate predominantly into secondary chloronemata, a minority differentiate into secondary caulonemata or gametophore buds. The radiating peripheral caulonemata tend to curve and secondary structures arise from them on the outside of the curve. Our observations of this pattern have confirmed most of the findings documented by McClelland (1987, 1988), Russell (1993) and Antonishyn (2024) and extended them with numerous details. Some of the differences between Antonishyn et al. and Raju and Ashton, e.g. the absence of predominance of clockwise over anticlockwise curvature of caulonemata growing over the surface of the solid medium, probably derived from the frequent positional changes with respect to the light source undergone by colonies in the latter study resulting from their removal from the growth cabinet for examination every two days. Possible mechanistic causes of caulonemal curvature in this fundamentally heterotrichous, morphogenetic system, which typically results in a spiral arrangement of the peripheral caulonemata, have been discussed in detail in Antonishyn et al. (2024) and will not be elaborated upon here.

The growth (increase in maximum width, w) curve of gametophytic colonies, mainly due to the centrifugal extension of primary and secondary caulonemata, is sigmoidal (**Fig. 6a**) and the instantaneous growth rate (dw/dt) increases until day 18 or 19 of our experiment before declining (**Fig. 6b**). This accords quite well with the discovery by Proust et al. (2011) that, in three week old *P.patens*, strigolactones originating from gametophores act as diffusible signals inhibiting caulonemal elongation and colony expansion. Differences in reported cell cycle times of caulonemal apical cells, 5.5 -10 h (McClelland 1987, Cove 1992, Vidali and Bezanilla 2012, Antonishyn et al. 2024) may reflect different growth rates at different times in the culture period. Thus, the increasing growth rate up to day 18 or 19 of our experiment may indicate that the caulonemal apical cell cycle time becomes progressively shorter before becoming longer under the influence of strigolactones. Alternatively, the differences may be attributable to differences in culture conditions, even when minor, that frequently and almost unavoidably occur in different laboratories.

### Most of the earliest gametophore buds are arranged in a fairy ring

We observed that the rate of production of gametophores (dn/dt) (**Fig. 6e**) accelerated maximally on day 12 (d^2^n/dt^2^) (**Fig. 6f**) and was highest on day 13 of the experiment, with a sharp peak in the rate between days 11 and 15 (**Fig. 6e**). The production rate declined sharply from day 13 until day 15 and thereafter decreased more slowly. Most of the gametophores that had formed between days 11 and 15 were arranged in a fairy ring in approximately three weeks old colonies. Considering a bud formed on day 13 and assuming an average caulonemal cell cycle with a duration of about 8 h and the differentiation of caulonemal apical cells from day five or six, we calculate that each caulonema comprises between 45 and 48 subapical cells with usually a single bud situated with very approximately 21-24 caulonemal subapical cells proximal to it and about 24 cells distal to it. In older colonies, the fairy ring is obscured by continued bud production, albeit at a slower rate, that is responsible for infill of the centre of the ring and formation of buds on more distal caulonemal subapical cells generated by further divisions of the apical cell of primary and secondary caulonemata. Infill of the centre of the fairy ring probably results from the differentiation of previously dormant, proximal SBIs or from the basal-most cell of secondary chloronemata. We have considered several possible mechanistic explanations of the coordinated formation of gametophore buds and their arrangement in a fairy ring. Below, we describe and discuss three of them.

### A completely cell autonomous model in which the fates of SBIs are determined stochastically by random probabilities

Based on McClelland’s observations, these probabilities would be approximately 85-90% for secondary chloronemata, 5-6% for secondary caulonemata, and <1% for buds. This explanation is appealing but one we have rejected, at least in its simplest form, for the following reasons. The probability of a SBI differentiating into a bud is greatly increased by buds developing on side branches adjacent to it on the same caulonema. As a result, buds tend to be clustered (Russell 1993, Cove et al. 2006, **Fig. 4a (panel C)**). Random probabilities of SBI fates are unlikely to generate bud clusters of this kind. Also, we have employed computer modelling using L-systems (Lindenmayer 1968a and b), to identify and assemble a set of production rules for the development of a virtual moss. Similar random probabilities to those listed above were used in the modelling process. Although we succeeded in generating a very realistic virtual moss gametophyte, its gametophores were never arranged in a fairy ring (Ashton et al. 1993, Fracchia & Ashton, 1995). Lastly, studies with *P. patens* mutants altered in their responses to auxin and/or cytokinin (Ashton et al. 1979, Cove and Ashton 1984) reveal that cytokinin is required for bud formation in an auxin-dependent manner. Drip-feeding experiments using liquid media with and without auxin and/or cytokinin confirm this (Cove and Ashton 1984). Thus, a completely cell autonomous stochastic mechanism is untenable and even a modified model incorporating, for example, stochastic variations in SBI sensitivity to cytokinin and/or auxin, does not circumvent the issues of bud clustering and the temporal and spatial organisation required to produce this morphogenetic pattern.

### A model in which the position (age) of the caulonemal subapical cell subtending a SBI determines its fate

The first SBIs are usually present on the second and/or third caulonemal subapical cells and their developmental fates are determined at this stage or only slightly later since the differentiation of derivative structures is often discernible from subapical cells in the fourth position. It seems improbable that the position of a subapical cell within this narrow positional window in a caulonema determines the fate of its SBI since each new subapical cell acquires the position (age) occupied by its predecessor one apical cell cycle earlier and it, in turn, acquires the position (age) occupied by its predecessor. Thus, each subapical cell occupies successively the same positions as its predecessors during the time frame when its fate is determined. Therefore, we have discounted this model also and suggest instead that it is the global age of the moss colony rather than the age of individual cells that is the key to understanding the temporally and spatially coordinated organisation of buds into a fairy ring.

### A model in which the global age of the moss colony fixes SBI fates

Gametophore buds begin to appear about 11 days after inoculation and the instantaneous rate of bud production (dn/dt) increases very sharply during days 11 to 13, declines steeply between day 13 and 15 and thereafter decreases more slowly (**Fig. 6e**). The sharp peak in dn/dt between days 11 and 15 corresponds to the time when fairy rings are formed. The auxin-dependent requirement of cytokinin for bud induction is well documented (Ashton et al. 1979, Cove and Ashton 1984) and we propose that the observed changes in the rate of bud production reflect earlier changes in the intracellular and/or extracellular levels of cytokinin. This model is strongly supported by the findings of Schulz et al. (2000), who showed that a peak in the concentration of isopentenyladenine (iP), the major cytokinin of *P. patens*, in the culture medium occurs nine days after inoculation and precedes the formation of buds on day 13. They suggested that the iP-maximum at day nine acts as the trigger for bud formation. This accords well with our observation that acceleration (d^2^n/dt^2^) of the rate of bud production increases soon after this on day 11 and reaches a maximum by day 12. A majority of cytokinin and of auxin is extracellular, i.e. in the medium, and both Reutter et al. (1998) and Schulz et al. (2000) proposed that communication between caulonemata might be mediated by diffusible morphogens including auxin and cytokinin. We suggest that fluctuation of the cytokinin level in the medium could be responsible for the temporally and spatially coordinated arrangement of buds in a fairy ring and for observed bud clustering on individual caulonemata. An appealing extension of this model could also explain the probabilities of the various developmental fates of SBIs. We argue, as has Cammarata et al. recently (2023), that it is not the absolute values of cytokinin and auxin that are the determinants of morphogenesis but rather the ratio of active cytokinin to active auxin. Thus, we envisage overlapping fluctuations in cytokinin and auxin in the medium during moss cultivation resulting in changing values of the cytokinin/auxin ratio with time. We postulate that a minimal value, or higher, of cytokinin/auxin is required for induction of budding and a maximal value of cytokinin/auxin, or lower, (i.e. a minimal value of auxin/cytokinin or higher) is needed for the formation of caulonemata, while all intervening values support the differentiation of SBIs into secondary chloronemata.

## Acknowledgements

*In memoriam*: Manjarabad V.S. Raju, Ph.D., a good colleague and friend, who died on March 31, 2012.

This work was supported in part by a grant from the Faculty of Graduate Studies and Research, University of Regina, Regina, Saskatchewan awarded to M.V.S. Raju.

